# Passive motor learning: Oculomotor adaptation in the absence of feedback on behavioral errors

**DOI:** 10.1101/2020.05.13.095018

**Authors:** Matan Cain, Mati Joshua

**Author notes:** Correspondence: Matan Cain, The Edmond and Lily Safra Center for Brain Sciences, The Hebrew University of Jerusalem, Edmond J. Safra Campus, Jerusalem 9190401.

## Abstract

Motor adaptation is commonly thought to be a trial-and-error process in which accuracy of movement improves with repetition of behavior. We challenged this view by testing whether erroneous movements are necessary for motor adaptation. In the eye movement system, the association between motor command and errors can be disentangled, since errors in the predicted stimulus trajectory can be perceived even without movements. We modified a smooth pursuit eye movement adaptation paradigm in which monkeys learn to make an eye movement that predicts an upcoming change in target direction. We trained monkeys to fixate on a target while covertly, an additional target initially moved in one direction and then changed direction after 250 ms. Monkeys showed a learned response to infrequent probe trials in which they were instructed to follow the moving target. Further experiments confirmed that probing learning or residual eye movement during fixation did not drive learning. These results show that movement is not necessary for motor adaptation and provide an animal model for studying how passive learning is implemented. The standard model assumes that the interaction between movement and error signals in the cerebellum underlies adaptive learning. Our results indicate that either sensory inputs are sufficient for driving learning in the cerebellum or that learning is implemented partly outside the cerebellum.

## Introduction

To better understand learning, signals that drive learning need to be identified behaviorally to reveal its implementation at the neuronal level. Motor adaptation is an especially valuable model for studying learning since experiments can reproducibly generate an experimental perturbation and then follow the changes in behavior on a trial-by-trial basis. Recent research has highlighted the importance of motor commands in driving motor adaptation. For example, the difference between the predicted and actual consequences of movement was shown to have both a computational advantage and account for behavioral results (Miall and Wolpert 1996; Shadmehr et al. 2010). However, motor command is only one of many signals that may drive motor adaptation (Mazzoni and Krakauer 2006; McDougle et al. 2016; Mostafa et al. 2019).

In terms of implementation, it has been hypothesized that in the cerebellum, movement and sensory signal converge to drive learning (Wolpert et al. 1998). When an erroneous motor command is executed, the climbing fiber input to the cerebellum drives plasticity that results in more accurate upcoming movement (Gilbert and Thach 1977; Ito 1982; Stone and Lisberger 1990). In the eye movement system, there is impressive trial-by-trial evidence for an association between climbing fiber input (manifested as complex spikes), the simple spike output of the Purkinje cell and learned behavioral changes (Herzfeld et al. 2018; Medina and Lisberger 2008; Suvrathan et al. 2016). In addition, in the eye movement system there is extensive data on which cerebellar sites drive eye movement and the pathways that provide signals to these areas (Voogd et al. 2012). Identifying non-motor signals in oculomotor adaptation can be interpreted in the context of what is already known about the implementation of motor learning and lead to testable hypothesis on where and how non-motor signals drive adaptation. Thus, animal models which use the eye movement system provide an excellent framework to determine how non-motor signals drive motor adaptation.

We modified a smooth pursuit eye movement leaning paradigm to test whether motor commands are needed for learning to occur. When monkeys (and humans) are trained to track a moving target that repeatedly undergoes the same change in direction at a predictable time, a learned smooth pursuit eye movement is elicited prior to change in target direction (Joshua and Lisberger 2012; Medina et al. 2005). These behavioral changes occur quickly and asymptotically after a few as 50 trials (Hall et al. 2018). During learning of perturbed target motion, the relationship between motor command and prediction target trajectory can thus be teased apart because motion can be sensed covertly without eye movement. We therefore designed a new paradigm in which monkeys learned to predict a change in direction of a target without tracking it. We termed this *passive motor learning.* We examined this type of learning in infrequent trials in which monkeys tracked a moving target, to show that monkeys can learn passively by observing and not tracking target motion.

Our findings raise questions and testable hypothesis as to the areas and mechanisms involved in passive learning. If passive learning involves the cerebellum, the current results suggest that known mechanisms for learning can be driven by sensory inputs alone. Alternatively, other areas might implicate the passive learning component. An intriguing possibility for learning is the frontal eye field-supplementary eye field (FEF-SEF) loop which uses visual cues to plan eye movements (Chen and Wise 1995a; Fukushima et al. 2011).

## Methods

We collected behavioral data from two male Macaca Fascicularis monkeys (4-6 kg). All procedures were approved in advance by the Institutional Animal Care and Use Committees of the Hebrew University of Jerusalem and were in strict compliance with the National Institutes of Health Guide for the Care and Use of Laboratory Animals. We implanted head holders to restrain the monkeys’ heads in the experiments. After the monkeys had recovered from surgery, they were trained to sit calmly in a primate chair (Crist Instruments) and consume liquid food rewards (baby food mixed with water and infant formula, 0.1 mL /trial) from a tube set in front of them. We trained the monkeys to track spots of light that moved across a video monitor placed in front of them.

### Visual stimuli and experimental design

Visual stimuli were displayed on a monitor 65 cm from the monkeys’ eyes. The stimuli appeared on a dark background in a dimly lit room. A computer performed all real-time operations and controlled the sequences of target motion. The position of the eye was measured with a high temporal resolution camera (1 KHz, Eye link - SR research) and collected for further analysis. Monkeys received a reward when tracking the target successfully.

We used eye velocity and acceleration thresholds to detect saccades automatically and then verified the automatic detection by visual inspection of the traces. The velocity and acceleration signals were obtained by digitally differentiating the position signal after we smoothed it with a Gaussian filter with a standard deviation of 5 ms. Saccades were defined as an eye acceleration exceeding 1000°/s^2^, an eye velocity crossing 15 °/s during fixation or eye velocity crossing 50°/s while the target moved. We first removed the saccades and treated them as missing data. We then averaged the traces with respect to the target motion onset. Finally, we smoothed the traces using a moving average filter with a span of 21 ms.

Pursuit stimuli were presented in trials. In eye movement trials, each trial started with a circular white target that appeared in the center of the screen. After 1s of presentation, in which the monkey was required to acquire fixation, the target stepped to a 4° eccentric position and started to move in the opposite direction at 20 °/s (step-ramp, Rashbass and Westheimer, 1961). After 250 ms, an orthogonal component of motion was added. The velocity of the orthogonal motion varied according to the type of eye movement trials. In test trials, the velocity of the added component was 20 °/s. In small angle trials, this velocity was 0.5 °/s. In probe trials, the target did not change direction. After additional 400 ms (650 ms after motion onset), the target stopped and stayed still for an additional 500 ms. The monkey was rewarded at the end of trials for keeping it eyes within a window around the target of 5×5° (3×3° before a change in direction). We used a large fixation window so that the monkeys’ behavior was not restricted during the learning trial. We prescreened the monkeys’ behavior to select target motion directions in which we could consistently drive learning. Specifically, these directions consisted of down and to the right for base and learning directions.

In fixation trials, two targets were displayed: a stationary and a moving target. The stationary target was a 1° side length square. In all the fixation trials, the stationary target was displayed during the entire trial. To be rewarded, the monkey had 400 ms to acquire fixation and had to keep its gaze within a 5×5° window throughout the entire trial (3×3° before a change in direction of the moving target). We used a large fixation window to match the fixation window in the learning trials. We verified the potential confound that the monkeys might initially track the moving target although they were instructed to fixate. We confirmed that the monkeys only made very small eye movements during the fixation trials and we designed experiments to control for this movement. The moving target was a white circular spot (except on reward blocks, see below), similar to the target on the eye movement trials. In fixation learning trials the moving target moved along the same trajectory as in eye movement trials. Fixation learning block comprised 90 fixation trials and 10 test or probe trials. Variants of this basic fixation learning block are described in the results section. Learning blocks were interleaved with washout blocks of 100 trials made up of 50% probe trials and 50% fixation trials in which the target moved but did not change direction.

The learned eye velocity was computed as the average eye velocity during the 100 ms around the change in direction in test trials minus the daily average eye velocity of the last 25 eye movement trials in the washout blocks. We adjusted the signs of the data such that positive values of learning indicate eye velocity in the learning direction. We estimated the growth of learning (L) over trials by fitting a sum of two exponentials to the learned responses.

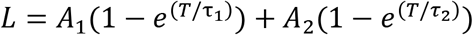

where A_x_ is the peak magnitude of learning, *τ_x_* is the “trial constant” of learning, and T+1 is the trial number.

## Results

### Learning to predict change in target direction by observation

We used a smooth pursuit eye movement learning paradigm in monkeys (Fig. 1A), to test whether feedback on behavioral errors is needed to adjust behavior. The first step consisted of a motor learning block (Joshua and Lisberger 2012; Medina et al. 2005) where the monkeys tracked a single moving target that changed direction 250 ms after the onset of motion (Fig. 1A top, *eye movement trial).* We term the direction in which the target initially moved the *base direction* (downward in Fig. 1A) and the orthogonal direction in which we later added a velocity component the *learned direction* (rightward in Fig. 1A). In the initial learning trials, the eye movement in the learned direction was reactive rather than predictive. After the target changed direction, the eye moved abruptly with a visually driven characteristic reaction time (about 100ms) takes place (Fig. 1B gray line). After several repetitions of trials with a change in direction the monkeys learned to predict the upcoming motion and moved their eyes in the learned direction even before the target changes direction (Fig. 1B black line, arrow points to the learned component). In this paradigm the predictive eye velocity was not sufficient to completely match the upcoming target motion, so that the monkeys still abruptly responded to the change in direction (Fig. 1B, black line) which was often followed by a catchup saccade (not shown). To avoid confounding the learned with the visually driven response, the analysis here was restricted to the first 300 ms after motion onset in the base direction (Fig. 1C).

**Figure 1.**
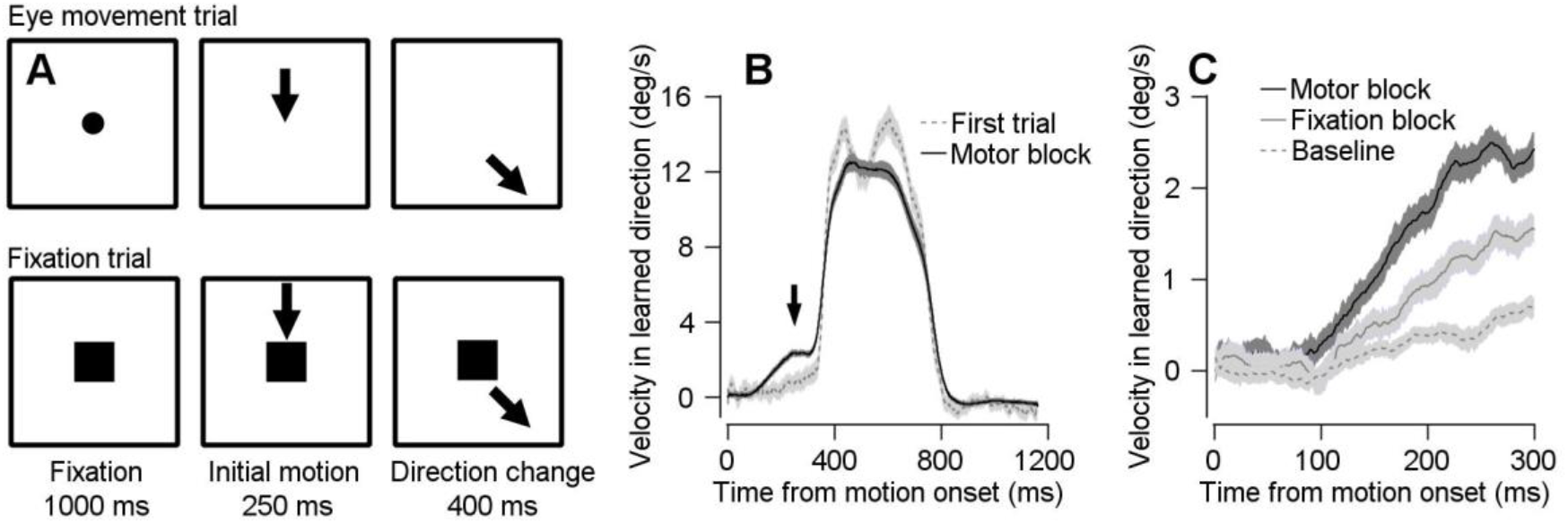
Trial schematics and behavior in motor and fixation blocks. **A:** Schematics of the eye movement (top) and fixation (bottom) trials. Arrows show the direction of target motion, circle represents the target prior to motion onset and squares represent the fixation target. **B:** Average eye movement in the learned direction in the first trial of learning (dashed gray trace) and post-learning trials on motor blocks (black). **C:** Average eye movement in the learned direction at the end of washout blocks (dashed gray) and after learning on eye movement trials on motor blocks (solid black) and fixation blocks (solid gray). In all traces, shadowing represents SEM.

Most theories of motor learning posit that sensory feedback on movement errors in learning trials and corrective behavior drive subsequent learning (Ito, 1972; Ito and Kano, 1982; Wolpert, 1998). To test whether the feedback on eye movement is necessary for learning, we designed an additional learning block, termed fixation block, in which the target changed direction, but the monkey did not follow it. In most trials (90%) the monkeys were required to maintain fixation at a square in the center of the screen while the moving target changed direction (Fig. 1A - bottom, *fixation trial).* Unlike the eye movement trials, in the fixation trials the monkeys were passive: the monkeys fixated the center of the screen, which prevented them from tracking the moving target and responding to the change in motion direction.

We tested learning in a small fraction of trials (10%) in which the square fixation target was not displayed, and the monkeys were required to follow the moving target exactly as in eye movement trials (Fig. 1A, top). In these trials, the monkeys shifted their gaze in the direction of motion even before the target changed direction (Fig. 1C, gray solid trace). To assess whether the monkeys indeed learned from these fixation trials we compared the learned response in the fixation blocks and the end of the washout blocks. The washout blocks consisted of 100 trials in which the target never changed direction (see Methods). By the end of the washout block (the last 50 trials, termed baseline trials), the eye velocity in the learning direction was close to zero (Fig. 1C, dashed trace). We quantified the learned response as the average eye velocity in the learned direction between 200 and 300 ms after motion onset. The learned response was maximal for the motor learning blocks, intermediate in the fixation blocks and the smallest in washout blocks (Friedman test, p =1*10^-12^, Post hoc signed rank test with Bonferroni correction, motor > fixation p = 1.2*10^-9^, fixation > washout, p=2.5*10^-9^, n= 46). Thus, in sessions with infrequent eye movement trials, the monkeys were still able to adjust their behavior to the change in target motion, suggesting that learning was acquired in fixation trials without movement.

### Movement in infrequent trials does not explain the learned response in fixation blocks

Next, we ruled out the possibility that learning in fixation blocks is driven solely by the infrequent trials (10%) in which the monkeys tracked the target. We tested the behavior of the monkeys in additional learning blocks in which the target did not change direction on the fixation trials (Fig. 2A). We termed these blocks *incongruent learning blocks* (Fig. 2A, right) and the block in which the target changed direction in fixation trials as it did in the movement trials *congruent learning blocks* (Fig. 2A, middle). Hence, the learned response in the fixation incongruent learning blocks could only result from the repetition of the eye movement trials. Thus, if learning was driven solely by infrequent eye movement trials, we would expect that the learned response should be similar on the congruent and incongruent blocks. When tested on the infrequent (10%) eye movement trials the eye velocity in the learned direction was lower in the incongruent versus the congruent learning blocks (Fig. 2B). Paired comparisons between nearby congruent and incongruent blocks that were recorded the same day (but separated by at least one washout block - see Methods) indicated that in most sessions, the learned response was higher in congruent blocks than in incongruent blocks (Fig. 2C, signed rank test p=5.9*10^-6^). These results indicate that fixation trials play an important role in the development of the learned response.

**Figure 2:**
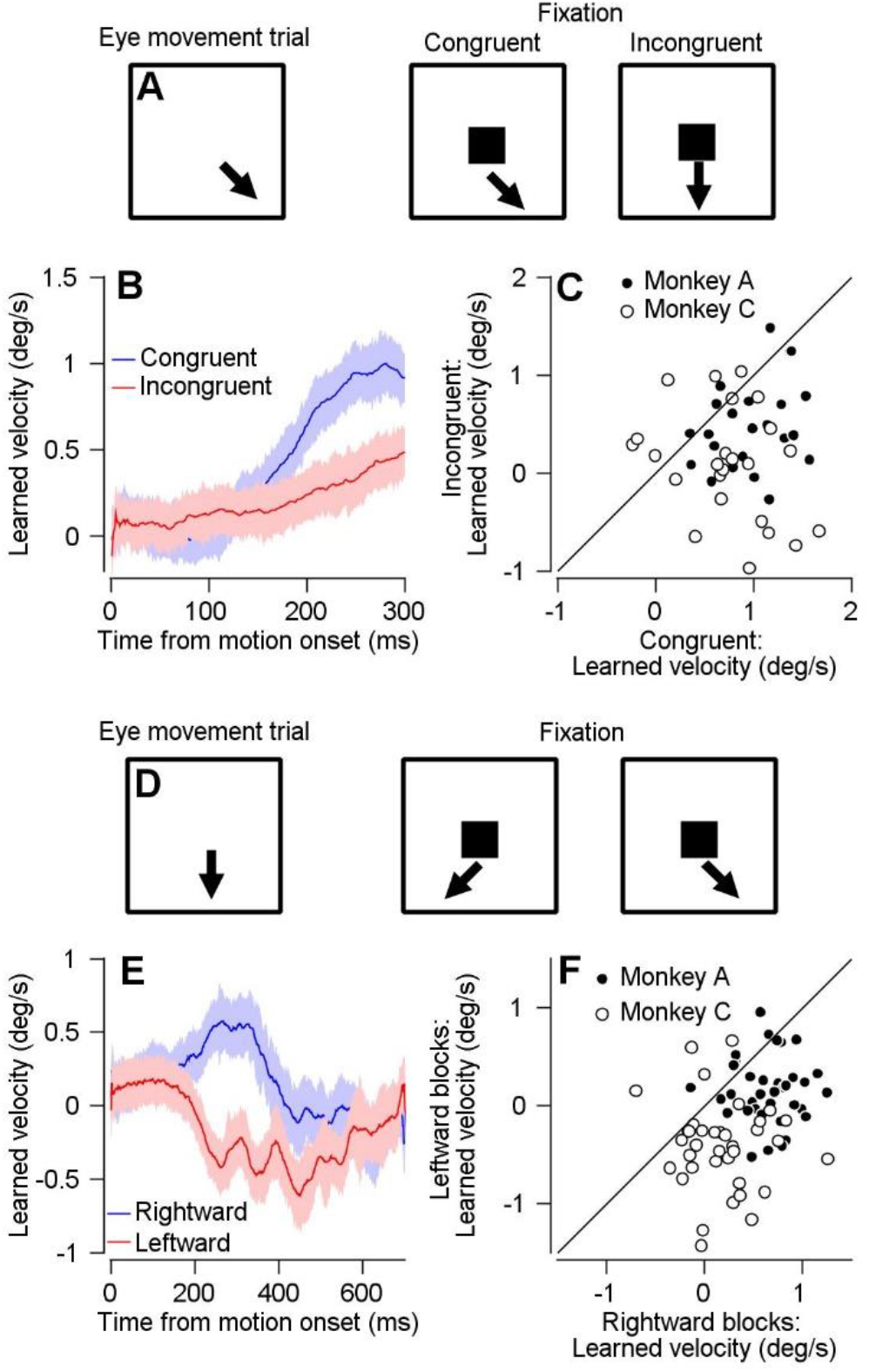
Learning from observation is not driven solely by infrequent eye movement trials. **A:** Schematics represent the direction change epoch in the different experimental conditions. Left: eye movement trials with change in direction, middle: congruent trial-fixation trial with directional change, Right: incongruent trial-fixation trial without directional change. **B:** Average learned eye velocity as a function of time from motion onset on congruent (blue) and incongruent (red) blocks. **C:** Learned eye velocity on incongruent (vertical) versus congruent (horizontal) blocks. Filled and open symbols show data from monkeys A and C. Solid line indicates unity. **D:** Schematics represent the target motion in the different experimental conditions. Left: eye movement trials without a change in direction, Middle: fixation trials in which rightward is the learned direction, Right: fixation trial in which leftward is the learned direction. **E:** Average learned eye velocity as a function of time from motion onset in blocks in which the target moved rightward (blue) or leftward (red). **F:** Learned eye velocity in adjacent blocks in which on fixation trials the target moved rightward (horizontal) or leftward (vertical). Filled and open symbols show data from monkeys A and C. Solid line indicates unity. In all traces, shadowing represents the SEM.

This conclusion relies on the assumption that the contribution of the eye movement trials to the learned response is identical in the fixation congruent and incongruent blocks. To further confirm that monkeys indeed learned from the congruent fixation trials we tested additional learning blocks. As in the fixation congruent trials, the target changed direction in the fixation trials but unlike the previous learning blocks we probed learning using trials in which the target did not change direction (these trials were thus identical to the baseline trials described above) (Fig. 2D, left). The only signal that could be used for learning in these blocks was the change in direction in the fixation trial. We alternated blocks in which the fixation trials had opposite learned directions, i.e., left (Fig. 2D, middle) or right (Fig. 2D, right). Thus, this experimental design had the advantage that in each learning block the monkeys never followed a target moving in the learned direction in eye movement trials and that on the fixation trials the target always changed direction.

In eye movement trials the average eye velocity deflects towards the learned direction (Fig. 2E). Positive and negative values in this analysis indicate movement right and left. Importantly, this deflection is not visually-driven because the stimulus in eye movement trials did not have any motion in the learned direction. Therefore, this deflection must result from learning in fixation trials alone. To directly compare sessions, we plotted the learned component in alternating blocks with opposite learned directions. The bias in the learned response towards the change in direction is manifested by the strong tendency of the dots to plot beneath the equality line in Figure 2F (signed rank test, p=7.7*10^-10^). We found a slight difference between monkeys. In Monkey C the bias was symmetric, i.e., in each learning block the eye moved towards the direction of the change in target motion (positive and negative horizontal and vertical values, in open dots in Fig. 2F). The movement of Monkey A was slightly biased towards positive values (corresponding to motion to the right), as indicated by the positive values on the horizontal axis and close to 0 on the vertical axis shown by the open dots in Fig. 2F. Nevertheless, the comparison between blocks indicated that in both monkeys the change in direction in fixation trials biased learned eye velocity in the corresponding direction. Thus, the monkeys learned from passive learning, when the only signal for learning was the change in target direction on the fixation trials.

### Control for movements in the fixation window

So far, we have shown that monkeys learn from fixation trials, suggesting that neither the corrective movement nor the feedback on erroneous behavior is necessary for learning. One possible confounding effect is that monkeys did not completely suppress behavior in fixation trials (solid traces in Fig. 3B and C). To control for this eventuality, we conducted experiments to confirm that the behavioral responses in fixation trials did not affect the learned response.

**Figure 3:**
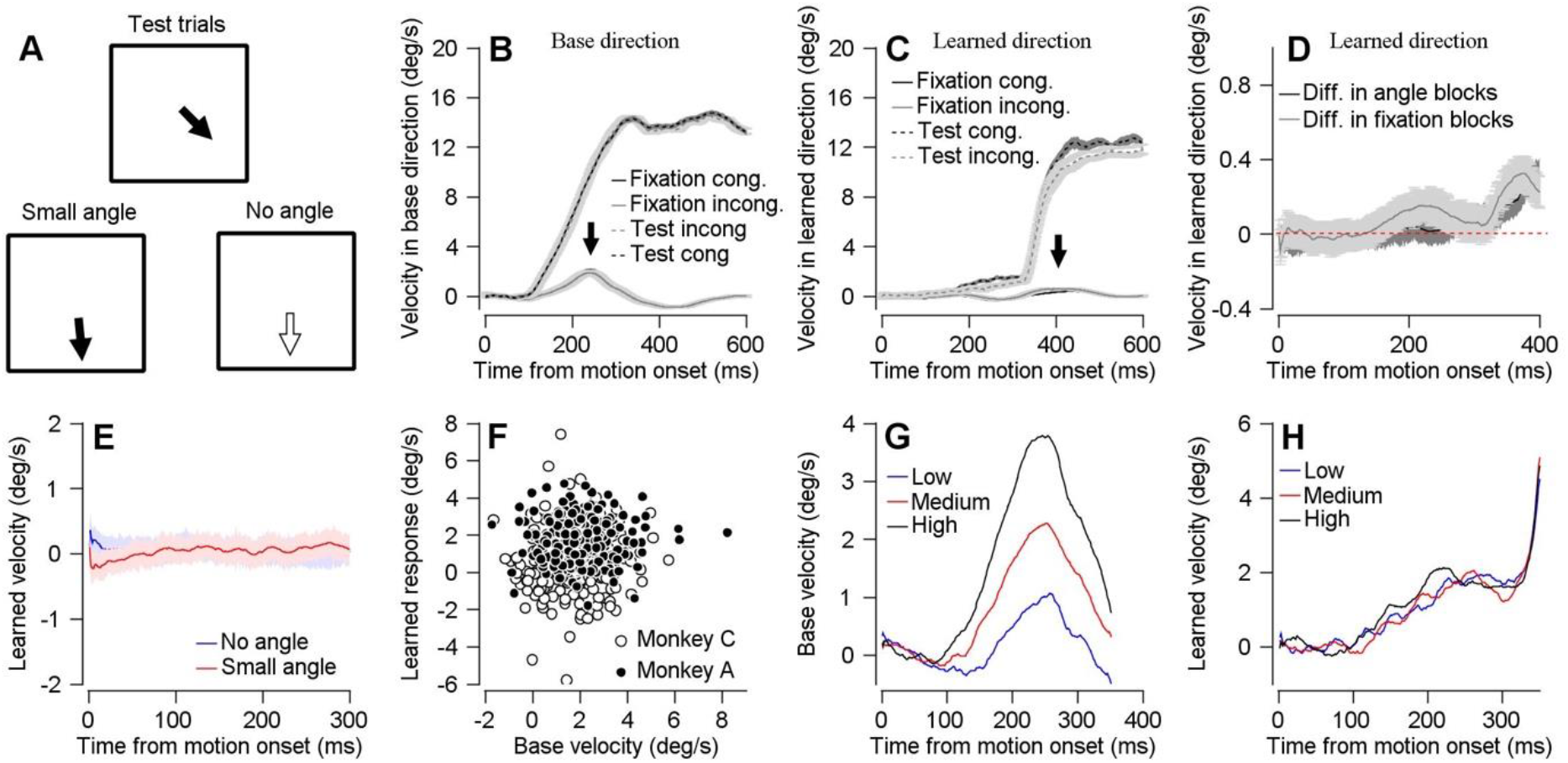
Learning is not driven by residual movement on fixation trials. **A:** Schematics showing the direction of motion change on trials with large (top), small (bottom left) and no change (bottom right) in target direction. **B, C:** Eye velocity in base (**B**) and learned (**C**) direction as a function of time from motion onset on fixation (solid trace) and test (dashed trace) trials. Black and gray traces show the velocity on the congruent and incongruent fixation trials **D:** Difference in learned eye velocity between fixation trials from congruent and incongruent blocks (gray) and small angle and no angle blocks (black). Dashed red line indicates null velocity **E:** Average learned eye velocity as a function of time from motion onset in blocks without change in direction (blue) and with a small change in direction (red). **F:** Base velocity on fixation trials, average from 200 ms up to 300 ms after motion onset versus learned response in subsequent test trials in congruent blocks. Filled and open symbols show data from monkeys A and C. **G, H:** Base velocity on fixation trials (**G**) and learned velocity on eye movement trials (**H**) as a function of time from motion onset for group of fixation trials with low, medium and high base velocities (blue, red and black traces). In all traces, shadowing represents the SEM.

In the learned direction in fixation trials, we observed a very slight increase in the velocity around the change in target direction in congruent trials compared to incongruent trials (arrow marking the dashed gray and black traces in Fig 3C and the gray trace in Fig. 3D). We aimed to mimic this behavioral difference to test whether it would impact the learned response on motor trials. To mimic the visually driven eye movement in the learned direction in congruent trials the monkeys were required in most trials (90%) to track a moving target that changed direction slightly after 250 ms such that a small component (0.5°/s) of the target velocity was added in the learned direction (Fig. 3A bottom left). In the second block, designed to mimic the behavior in incongruent trials, in 90% of the trials the target did not change direction (as in baseline trials described above, Fig. 3A bottom right). As expected, the difference in eye velocity in the learning direction between learning trials consisting of no angle and small angle blocks (Fig. 3D, black) was indeed similar to the difference between fixations trials in the congruent and incongruent blocks (Fig. 3D, gray). To keep the structures of the blocks as similar as possible and to probe learning, in the remaining 10% of the trials, the target changed direction as in the previous experiments (20°/s component in the learning direction, Fig. 3A top).

If indeed the corrective behavior we observed on the fixation trials was sufficient to drive learning we would expect to find a difference between the mimic blocks with and without the small angle. However, we found that the difference between the learned eye velocity on blocks with small and no angle was not significant (Fig. 3E, Wilcoxon signed rank test, p=0.26). Furthermore, the difference between the learned response in the congruent versus incongruent blocks was larger than the difference between blocks with and without an angle (ranked-sum p = 0.036). Therefore, this control suggests that the slight corrective movement we observed in the fixation trials does not drive learning.

Next, we focused on the increase in base velocity on fixation trials around the change in direction in both the congruent and incongruent blocks (solid gray versus black traces in Fig 3B, marked by an arrow). This movement might contribute to learning since the discrepancy between the movement and the direction of target change could elicit an error signal. However, if indeed this discrepancy between behavior and target motion drove learning, we would expect that larger movements in the base direction would correlate with more learning on the movement trials. However, in the congruent blocks, there was no significant correlation between base velocity averaged across the fixation trials and the amplitude of the learned response in the subsequent test trial (Fig 3F, the multiple regression analysis with monkeys and base velocity as predictors of learned velocity was significant for monkeys, p=3.02*10^-13^, but not for base velocity, p=0.34). Figure 3G and 3H demonstrate the absence of correlation in time for monkey A. We clustered the base velocity on the fixation trials into three groups according to the magnitude of the base direction eye velocity on the fixation trials (Fig. 3G). As expected from a non-correlated relationship, these clusters were not preserved when we plotted the learned response on the fixation trials (Fig. 3H). These result are consistent with previous research in the motor learning paradigm that did not find correlation between movement speed in the base direction prior to change in the target direction and learning in the next trial (Herzfeld et al. 2020). Thus, it is unlikely that residual movement on the fixation trials within the fixation window is necessary for learning.

### Learning in fixation blocks is driven by change in direction

We have shown that the monkeys were able to learn from fixation trials. We next attempted to better understand which component in the fixation trials was necessary for learning. In eye movement trials, the crucial instructive signal for learning is the change in target direction (Medina and Lisberger 2008; Yang and Lisberger 2014). Consequently, we tested whether motion in the learned direction of the target is essential to develop the learned response. Alternatively, information about the end point position of the target could be sufficient to drive learning. To answer this question, we compared the learned response in two learning blocks. The first block was identical to the fixation congruent block described above (Fig. 4A, top and middle). In this context we termed this block the *motion block.* The second block, termed the *position block* was similar to the previous block except that the moving target vanished right before the addition of the upward velocity component, 250 ms after motion onset. The target then reappeared at the end of the trial (650 ms after motion onset) in the same position as in the motion trials (Fig. 4A, bottom). We found that the learned response on the motion block was higher than on the position block (Fig. 4B). Single session comparisons indicated that this difference was significant (Fig. 4C, Wilcoxon signed Rank Test, p=1.8*10^-4^), consistent across monkeys and observed in most sessions. We conclude that the motion on the fixation congruent blocks plays an essential role in the development of the learned response.

**Figure 4:**
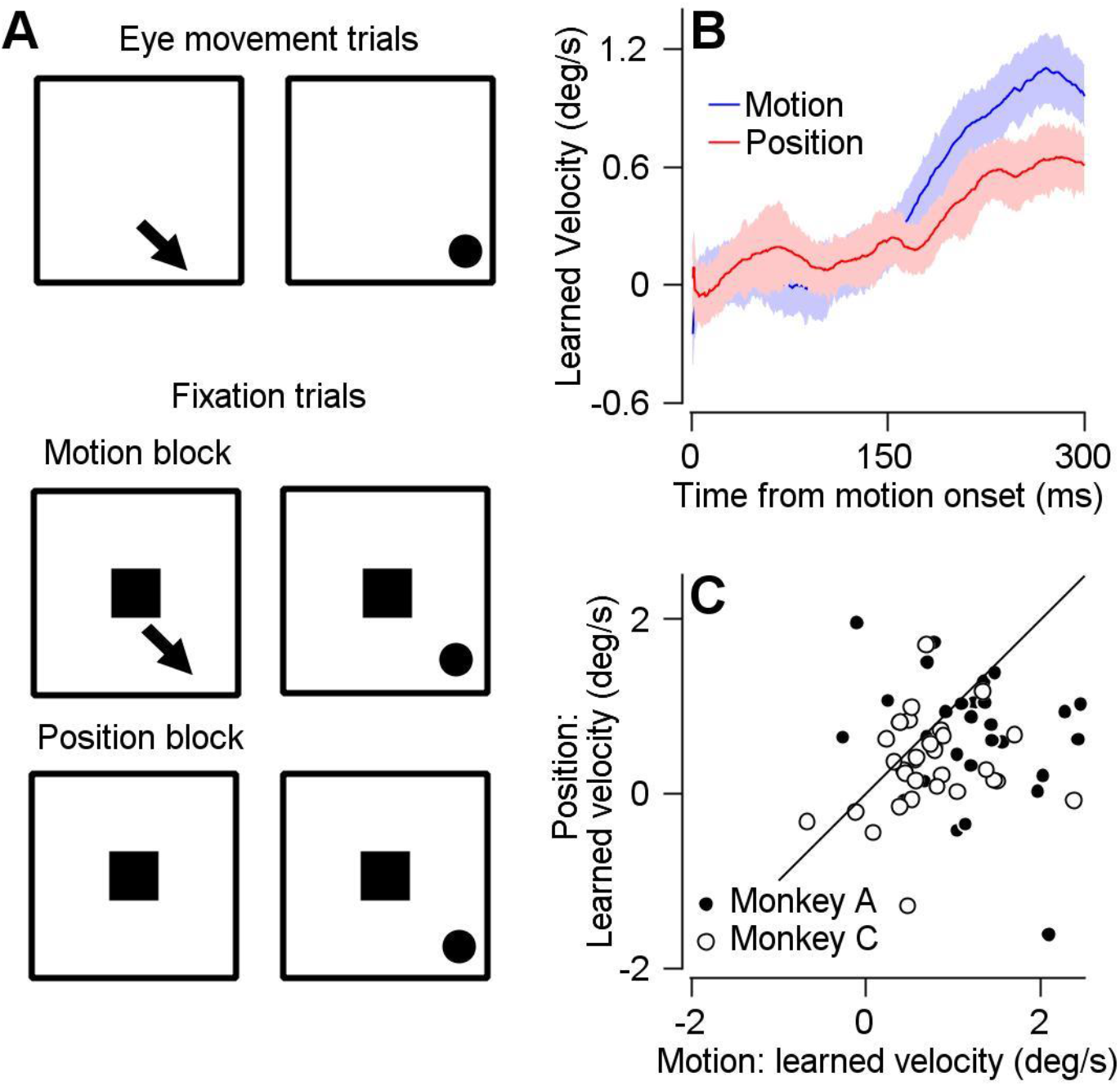
_Learning on fixation blocks is driven by change in direction. **A:** Schematics represent the target motion and position at the beginning and end of direction change epochs. Arrow represents the direction of motion; square represents the fixation target and dot represents the location of the moving target at the end of the trial. Top: eye movement trials with change in direction. Middle: Fixation trials in blocks with target motion. Bottom: Fixation trials in blocks without target motion - the moving target vanished with the change in direction and reappeared at the end of the epoch. **B:** Average learned eye velocity as a function of time from motion onset in learned direction in motion (blue) and position (red) blocks. **C:** Learned eye velocity on motion (horizontal) versus position (vertical) blocks. Solid line indicates unity. Filled and open symbols show data from monkeys A and C. In all traces, shadowing represents the SEM.

### Learned response in fixation blocks is modulated by expected reward

We have shown how basic sensorimotor parameters such as target motion and eye movements impact learning. We next tested whether the task’s broader context could also influence learning from observation. Specifically, we have previously shown that reward interacts with the visuomotor processing of the pursuit system (Joshua and Lisberger 2012; Lixenberg and Joshua 2018). We therefore designed a task to test whether the learned response could be modulated by reward information. The structure of the eye movement and fixation trials were similar to those described in the first part of the experiment (Fig. 1A). Each block consisted of 10% eye movement trials and 90% fixation trials. The fixation trials were equally divided (45%) into congruent trials and incongruent trials. The key difference was that the reward associated with each fixation trial was swapped between blocks. In the congruent-reward blocks, a reward was only given after congruent trials (Fig. 5A, top) whereas in the incongruent-reward blocks a reward was only given after incongruent trials (Fig. 5A, bottom). The color of the target indicated whether the monkey would be rewarded at the end of the trial (Fig. 5A). We used both rewarded and unrewarded trials in the same block to equalize the overall reward on each block to control for overall motivation changes. We tested learning in eye movement trials with a white target, in which the monkey always received a reward to ensure that the reward in these trials did not affect the expression of learning differently (Joshua and Lisberger 2012).

**Figure 5:**
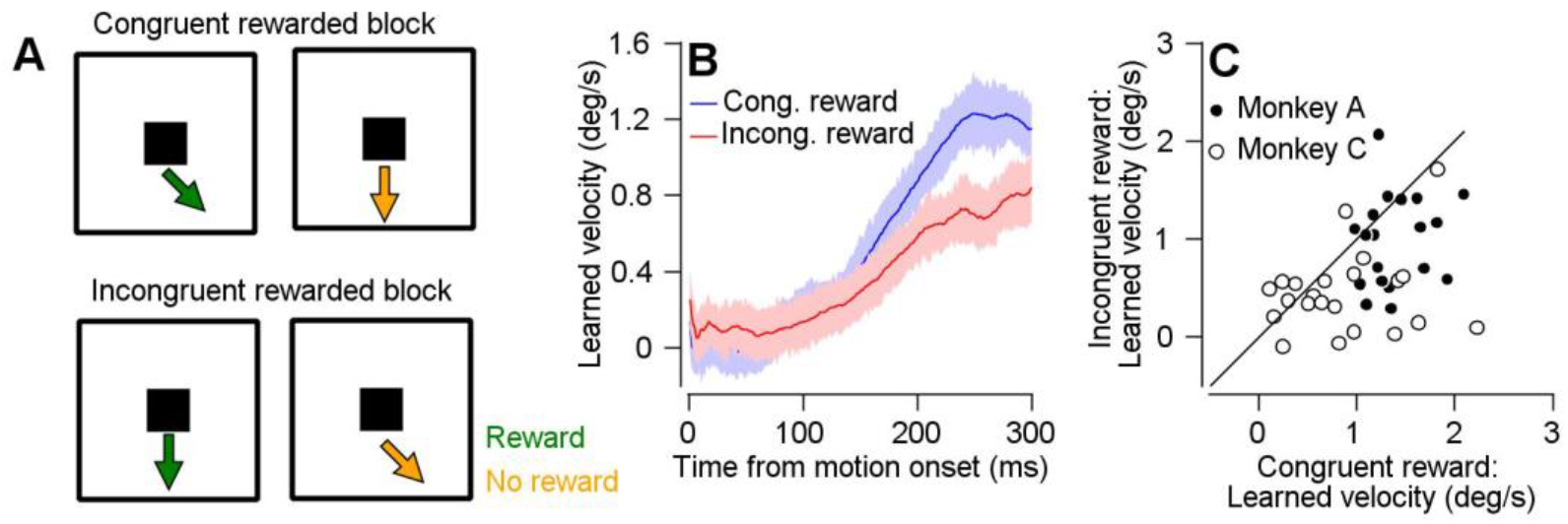
Learning on fixation blocks is modulated by expected reward. **A:** Top: fixation trials in congruent rewarded blocks-Left: congruent rewarded trials, Right: incongruent unrewarded trials. Bottom: fixation trials in incongruent rewarded blocks-Left: incongruent rewarded trials, Right: congruent unrewarded trials. **Colors correspond to the color of the target we used for monkey C. For monkey B we used blue and pink to signal reward and omission of reward. B:** Average eye velocity in post-learning eye movement trials as a function of time from motion onset in learned direction in congruent rewarded (blue) and incongruent rewarded (red) blocks. **C:** Eye velocity in congruent rewarded (horizontal) versus incongruent rewarded (vertical) blocks. Solid line indicates unity. One outlier that had values of (0.52; −1.53) is not shown. Filled and open symbols show data from monkeys A and C. In all traces, shadowing represents the SEM.

We found that reward modulated the amplitude of the learned response. The average learned eye velocity on the eye movement trials was higher for the congruent-reward than for the incongruent-reward blocks (Fig. 5B). Paired tests between interleaved blocks that were separated by a washout block indicated this difference was significant (p=6.1*10^-5^, signed rank Test) (Fig. 5C). These results corroborate the hypothesis that reward modulation affects the acquisition of learning, unlike previous findings in a motor learning context (Joshua and Lisberger 2012). This discrepancy might be due to differences between paradigms such as timing of the reward signal, direction of motion in reward versus non-rewarded trials or differences between learning on fixation and motor trials (but see Damasse et al. 2018; Liu et al. 2019). Note that prior to the experiment the monkeys were extensively trained to associate the color and the reward. Therefore, it is likely that the expected reward, rather than reward delivery, was the critical reward signal modulating learning, perhaps through attention mechanisms.

### Very rapid learning is probably explained by the uniformity of the learning block

In the previous sections we considered learning blocks as a whole without addressing the dynamics of learning. We were unable to reliably follow or quantify the learning curve of the observed learned response (Fig. 6B, dashed line), since most of the learning accrued prior to the first eye movement trial. In the fixation blocks, the learned eye velocity on the first eye movement trial (which was followed on average by 5 fixation trials) was not significantly different from the other trials (Fig. 6A, p=0.8, ranked-sum test). To explore the possibility that the lack of dynamics in learning in the experiment was due to very fast learning, we analyzed the learning curve in the interleaved motor learning blocks. We found that in these blocks most of the learning occurred very rapidly (Fig. 6B, solid). To quantify, we fit the learning curve to a double exponent (see Methods). We found that the rapid learning (*τ*_1_ = 4*10^-2^ trials) dominated the learning process in that it explained 68.46% of the learning in the first 100 trials.

**Figure 6:**
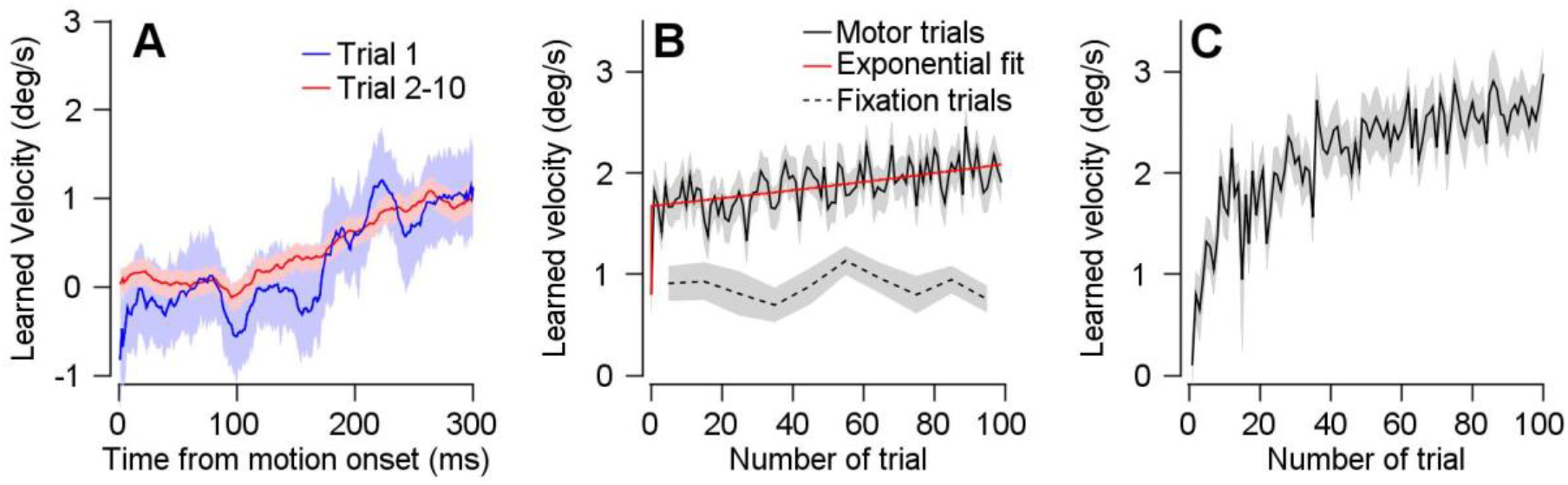
Very rapid learning may be accounted for by the uniformity of the learning block. **A:** Average eye velocity across sessions on eye movement trials in fixation congruent blocks for the first eye movement trial (blue) and remaining eye movement trials (red) **B**: Learning curve on motor (solid) and congruent fixation blocks (dashed) with fixed base and learned directions. Exponential fit of the motor learning curve is shown in red**. C:** Learning curve of motor blocks with change in the base and learned directions. In all traces, shadowing represents the SEM.

This fast learning was probably a result of the uniformity of the current experiment relative to previous experimental designs. We focused most of our experiments on one particular base and learning direction. We compared the learning curve during the eye movement trials to the learning curve from learning sessions in which we varied the base (0°,90 °,180 ° or 270°) and learning (clockwise and counter clockwise) direction. The learning curve in this richer context increased gradually (Fig. 6C). Thus, the speed of learning depended on the richness of the experimental context. Generally, other parameters such as the time between consecutive learning trials or different trials interleaved between learning trials could also influence the speed of learning.

## Discussion

### Passive motor learning

It is well-established that monkeys learn to predict a change in target direction when actively tracking the target (Medina et al. 2005). Here, we found that a passive observation of the change in target direction without tracking is sufficient to elicit a learned response. Thus, an association between motor output and sensory feedback is not necessary to elicit an adaptive response. In several other adaptation paradigms, sensory input alone was insufficient to elicit learning (Held and Freedr 1963; Mazzoni and Krakauer 2006; but see Mostafa et al. 2019). All these paradigms report a discrepancy between the predicted and observed sensory consequences of motor commands (Shadmehr et al. 2010). For example, application of a force field is known to change the observed sensory consequences of a given motor command. The smooth pursuit paradigm presented here differs from these paradigms in that the perturbation (the change in direction of the moving target) can be perceived without movement so that learning does not depend on the ability to predict the sensory outcomes of a motor command. This difference may explain why passive motor learning is possible in the smooth pursuit paradigm and highlights the important role of the oculomotor system in the study of motor adaptation driven by a sensory signal (in addition to sensory feedback).

The passive learning we found in monkeys could partially overlap with explicit motor learning in humans (Mazzoni and Krakauer 2006; McDougle et al. 2015, 2016) since neither require movement. Nevertheless, in pursuit, passive and motor learning of the sensory specificities of the instruction are important. We found that instructing learning without target motion was less effective in driving passive learning (Fig. 4). Furthermore, in pursuit motor learning, the monkeys relied on trial-by-trial fluctuations in that target motion direction rather than abstract rules such as alternation of the direction of motion (Yang and Lisberger 2010). Thus, passive (as well as motor) learning in smooth pursuit in monkeys is probably implemented through the sensorimotor representation of the target motion rather than an abstract explicit representation. The smooth pursuit eye movement system has been widely used as a model system for studying sensorimotor transformation and motor learning at the implementation level of neurons and networks (Joshua and Lisberger 2015; Lisberger 2010). The paradigm we developed here can be harnessed to provide testable hypothesis on where and how the brain implements passive learning. Another advantage of the paradigm stems from the temporal gap between the sensory inputs in fixation trials and their effect on later motor trials. Thus, this paradigm provides an easy way to dissociate between processing of visual motion and the generation of pursuit motor command.

### Possible Neural implementation in the cerebellum and frontal cortex

The cerebellar flocculus plays an important role in the development of a predictive response to an instructive change in target direction when subjects actively track the target during learning (Medina and Lisberger 2008). According to the classic cerebellar model, sensory errors resulting from inaccurate movement drive climbing fiber input (Albus 1971; Gilbert and Thach 1977). The climbing fiber input paired with input to the Purkinje cell results in an associative reduction in synaptic strength (Ito et al. 1982; Suvrathan et al. 2016). This reduction is thought to underlie the subsequent improvement in behavior.

It is possible that the same cerebellar mechanisms drive active and passive learning. At the neuronal level, this hypothesis predicts that all the hallmarks of cerebellar learning will be observed during passive learning. For example, in fixation trials climbing fiber inputs will be modulated during the target change of direction. Furthermore, the Purkinje cell simple spikes are likely to be tightly related to the climbing fiber input on a trial-by-trial basis (Herzfeld et al. 2018; Medina and Lisberger 2008; Suvrathan et al. 2016). The presence of a climbing fiber response after the change in direction on one trial should be associated with a change in the simple-spike firing rate on the subsequent fixation or eye movement trial.

Passive learning could be implemented in the FEF. Visual, motor and temporal signals converge in the FEF (Bruce and Goldberg 1985; MacAvoy et al. 1991; Schafer and Moore 2011; Schall et al. 1995; Schoppik et al. 2008; Sommer and Wurtz 2006). During pursuit learning neurons that are temporally tuned to the time of target change in direction are those that undergo the largest learning modulation (Li et al. 2011). If time tuning is preserved during fixation trials it might underlie passive learning. For example, during fixation, neurons that are tuned to the direction and time of the change in the target direction would respond the most vigorously. Any inputs to these cells from other cells that are tuned to the base direction prior to the time of change in direction would be potentiated through spike-timing dependent plasticity. This plasticity process should result in an increase in activity of neurons tuned to the learning direction even before the change in direction in fixation and motor trials.

Another possible learning mechanism would occur upstream from the FEF. The SEF is a good candidate for learning the association between the movement in the base direction and the addition of a component in a learned direction (Chen and Wise 1995b; Fukushima et al. 2004). The change in SEF activity would elicit a learned response through the reciprocal connections between SEF and FEF (Huerta et al. 1987). Thus, we can identify several plausible sites in which observed information is use to drive learning. Future work, probing and manipulating these networks, could use the paradigm we have established to study the implementation of motor adaptation without motor commands.

### Quantification of learning from fixation trials

The learned response shown in fixation blocks (Fig. 1) can be divided into two components: the passive learning elicited by fixation trials and the motor learning that resulted from the test trials. The trials assessing learning are also involved in the learning process; therefore, we cannot directly measure the learning elicited exclusively by passive learning. Indirect measures suggest that most learning in fixation blocks is due to passive learning. The learned response in the first test trial, which was proceeded only by fixation trials, was similar to the learned response late in learning (Fig. 6A) and the learned response in the incongruent blocks was small (Fig. 2B).

Although we cannot completely control for the magnitude of learning from eye movement trials, we can bound the amplitude of the learned response elicited by the fixation trials. The learned response in fixation blocks is an upper bound for the amplitude of the learned response elicited by fixation trials because it contains both passive learning and a small component of motor learning. The learned response in the experiment in which the target only changed direction on fixation trials (Fig. 2E) is a lower bound for the learning from fixation trials. In these blocks learning was assessed using non-adaptive probe trials that reduce the learning elicited by fixation trials. We quantified these bounds by calculating the ratio of the learned response in motor block to the learned response in the corresponding block. We estimated that passive learning in the current paradigm lay within a range of 18% and 48% (see Methods) of the total motor learning (learning in eye movement blocks; e.g., Fig. 1C). This estimation may not be the theoretical limit as it is possible that other non-motor factors could account for the difference between passive and motor learning. For example, attention or the exact location of the stimulus on the retina at the time of the direction change could have varied in eye movement and fixation trials. Further research should consider the interaction between learning mechanisms elicited by motor and non-motor signals in the presence of motor commands. Passive learning might be elicited concurrently with mechanisms driven by motor signals or alternatively, be elicited exclusively in the absence of motor signal.

Overall, we showed that the passive observation of target motion can drive behavior that was previously characterized as motor adaptation. We conducted controls and explored the conditions in which passive learning is expressed. The pursuit system provides a unique model system for studying passive learning since it can be explored at the implementation level in monkeys. We suggest possible mechanisms based on the known properties of smooth pursuit system. These hypotheses can become the basis for further investigations of passive motor learning in the pursuit and other systems.

## Acknowledgments

We thank Y. Botschko for technical assistance. This study was supported by a HFSP career development award Mati Joshua the Israel Science Foundation, and the European Research Council.

